# Statistical Methods for Detecting Circadian Rhythmicity and Differential Circadian Patterns with Repeated Measurement in Transcriptomic Applications

**DOI:** 10.1101/2022.06.05.494875

**Authors:** Haocheng Ding, Lingsong Meng, Chengguo Xing, Karyn A. Esser, Zhiguang Huo

**Affiliations:** Department of Biostatistics, University of Florida; Department of Medicinal Chemistry, University of Florida; Department of Physiology and Functional Genomics, University of Florida

## Abstract

Circadian analysis via transcriptomic data has been successful in revealing the clock output changes underlying many diseases and physiological processes. Repeated measurement design in a circadian study is prevalent, in which the same subject is repeatedly measured over time. Several methods are currently available to perform circadian analysis, however, none of them take advantage of the repeated measurement design. And ignoring the within-subject correlation from the repeated measurement could result in lower statistical power. To address this issue, we developed linear mixed model based methods to detect (i) circadian rhythmicity (i.e., Rpt_rhythmicity) and (ii) differential circadian patterns comparing two experimental conditions (i.e., Rpt_diff). Our model includes a subject-specific random effect, which will account for the within-subject correlation. Via simulations, we showed our method not only could control the type I error rate around the nominal level, but also achieve higher statistical power compared to other methods that cannot model repeated measurement. The superior performance of Rpt_rhythmicity and Rpt_diff were also demonstrated in two real data applications, including a human restricted feeding data and a human sleep restriction data. An R package for our methods is publicly available on GitHub to promote the application of our methods.

## 1 Introduction

Circadian rhythms are endogenous ∼ 24 hours cycles of behavior and physiology that have evolved as adaptions to the dark and light cycles caused by the earth’s rotation. Some examples of circadian behavior and physiology include sleep-wake cycles, body temperature, and melatonin secretion (Badia et al., 1991; Jung et al., 2010; Cagnacci et al., 1992; DIJK et al., 1992). In humans, the underlying circadian clock is found in virtually all cells throughout the body, with a master clock in the hypothalamic suprachiasmatic nucleus acting as a pacemaker to coordinate and synchronize the peripheral clocks (Hastings et al., 2003). The molecular mechanism underlying the circadian clock is a transcriptional/translational feedback loop driven by a set of core clock genes (Takahashi, 2017), including *CLOCK, BMAL1* as major transcriptional activators; and period family genes (i.e., *PER1, PER2, PER3*) and cryptochrome family genes (i.e., *CRY1, CRY2*) as major inhibitors. With the advancement of high-throughput sequencing technology, genomic insights into circadian clocks have recently come into focus (Koike et al., 2012), especially after 2017 when the Nobel Prize in Physiology or Medicine was awarded for the discoveries of molecular mechanisms controlling the circadian rhythm. In the literature, circadian transcriptomic landscape has been studied in many tissues including the postmortem brain (Chen et al., 2016; Seney et al., 2019), skeletal muscle (Hodge et al., 2015), liver (Hughes et al., 2009), and blood (Möller-Levet et al., 2013). Pan-tissue transcriptomic circadian analyses from humans (Ruben et al., 2018), mice (Zhang et al., 2014), and baboons (Mure et al., 2018) have revealed tissue-specific circadian patterns in gene expression. Beyond transcriptomics, circadian analysis was also performed in other types of omics data including DNA methylation (Lim et al., 2014), ChIP-Seq (Koike et al., 2012), proteomics (Wang et al., 2018), and metabolomics (Dallmann et al., 2012).

In the literature, several algorithms have been developed to detect circadian rhythmicity. The cosinor-based model assumes the expression level of a gene is a sine or cosine function of the circadian time (Cornelissen, 2014), which is biologically interpretable and enjoys accurate statistical inferences under Gaussian assumptions (Ding et al., 2021). Other parametric models, including Lomb-Scargle periodograms (Glynn et al., 2006) and COSOPT (Straume, 2004), can detect oscillating genes with irregular shape by assuming mixtures of sine/cosine curves with distinct frequencies. Nonparametric models, including ARSER (Yang and Su, 2010), RAIN (Thaben and Westermark, 2014), and JTK CYCLE (Hughes et al., 2010), are free of modeling assumptions, and thus also powerful in capturing irregular curve shapes. MetaCycle (Wu et al., 2016) aims to combine results from ARSER, JTK CYCLE and Lomb-Scargle using meta-analysis via the Fisher’s method. Both parametric and nonparametric algorithms have been widely adopted in circadian transcriptomic studies, and several review studies have systematically compared the performance of these algorithms (Hughes et al., 2017; Mei et al., 2020; Laloum and Robinson-Rechavi, 2020).

Though existing methods are promising to detect circadian rhythmicity, none of them is designed to handle the repeatedly measured data in a circadian experiment – multiple measurement of the same subject at different circadian time. For example, Lundell et al. (2020) recruited eleven overweight or obese men to examine the impact of restricted feeding on circadian transcriptome of skeletal muscle, with each participant being repeatedly measured every 4 hours over a 24-hours cycle. This type of circadian experiment with repeated measurement is prevalent in the literature (Perrin et al., 2018; Möller-Levet et al., 2013; Blume et al., 2017; Saner et al., 2021). These repeated measurement from the same subject are naturally correlated, and failing to consider these correlation structures may result in poor statistical powers in detecting circadian rhythmicity. To our knowledge, there are currently no existing methods that take into account the repeated measurement in circadian rhythmicity detection, and there is an urgent need for the circadian community to develop such toolboxes.

Another important research question in the circadian field is to detect differential circadian patterns underlying different experimental conditions (Hughey and Butte, 2016; Hsu and Harmer, 2012; Möller-Levet et al., 2013). Several epidemiology and animal studies have linked the disruption in clock and circadian gene expression with diseases including cancer (Sancar and Van Gelder, 2021; Ballesta et al., 2017), type 2 diabetes (Stenvers et al., 2019), sleep disorder (Möller-Levet et al., 2013), major depression disorder (Li et al., 2013), aging (Chen et al., 2016), schizophrenia (Seney et al., 2019), and Alzheimer’s disease (Lim et al., 2017). In the literature, all existing methods for detecting differential circadian patterns are based on the cosinor model, which can be categorized into two major types. The first type is to examine whether there is a difference in the overall circadian pattern, with the null hypothesis being the sine/cosine curve fittings are identical across two experimental conditions. Representative statistical methods of this first type include RobustDODR (Thaben and Westermark, 2016), HANOVA (Thaben and Westermark, 2016) and LimoRhyde (Singer and Hughey, 2019). The second type is to further pinpoint the exact difference — whether it is a change in amplitude, phase, MESOR (Midline Estimating Statistic of Rhythm), or goodness of fitness. And representative statistical methods of this second type include permutation approach (Chen et al., 2016), circaCompare (Parsons et al., 2020), and diffCircadian (Ding et al., 2021). However, these methods are not designed for repeated measured study. To our knowledge, there are currently no existing statistical methods that can handle repeated measurement data in differential circadian analysis, which consequently will result in a lack of statistical power in differential circadian analysis.

To close these research gaps, we develop a series of statistical methods based on linear mixed models in detecting circadian rhythmicity and differential circadian pattern (overall circadian pattern, the first type) with repeated measurement. We include the subject-specific random effect to model the repeated measurement from the same participant to increase statistical powers. The statistical inference is performed using the likelihood ratio test and the F test. We systematically evaluate the type I error and power of our proposed methods, and demonstrate their superior performance over existing competing methods (without considering repeated measurement) in both simulations and real data applications. The contribution and novelty of our proposed methods include: (i) to our knowledge, this is the first statistical method for detecting circadian rhythmicity in circadian experiment with repeated measurement; (ii) this is the first statistical method for detecting differential circadian patterns (overall circadian pattern; the first type) in circadian experiment with repeated measurement; (iii) we developed an R package for our proposed method, which is publicly available on GitHub, to promote its applications in the circadian field.

## 2 Method

To take advantage of the repeated measurement data structure, we developed linear mixed models for (i) circadian rhythmicity detection within one experimental condition and (ii) differential circadian analysis comparing two experimental conditions. The statistical inferences of these methods are based on the likelihood ratio test as well as the F test.

### 2.1 Notation for the sine curve fitting

As shown in Figure 1a, we assume the expression level of a gene *y* is a sinusoidal function of the circadian time, which follows the notation in Ding et al. (2021). To be specific, *A* is the amplitude; *M* is the MESOR (Midline Estimating Statistic of Rhythm); 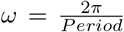 is the frequency. We set *period* = 24 hours to be consistent with the 24h diurnal cycle. *ϕ* is the phase of the sine curve. For the ease of discussion, we leave out the unit ‘hours’ in period, phase, and other related quantities.

**Figure 1:**
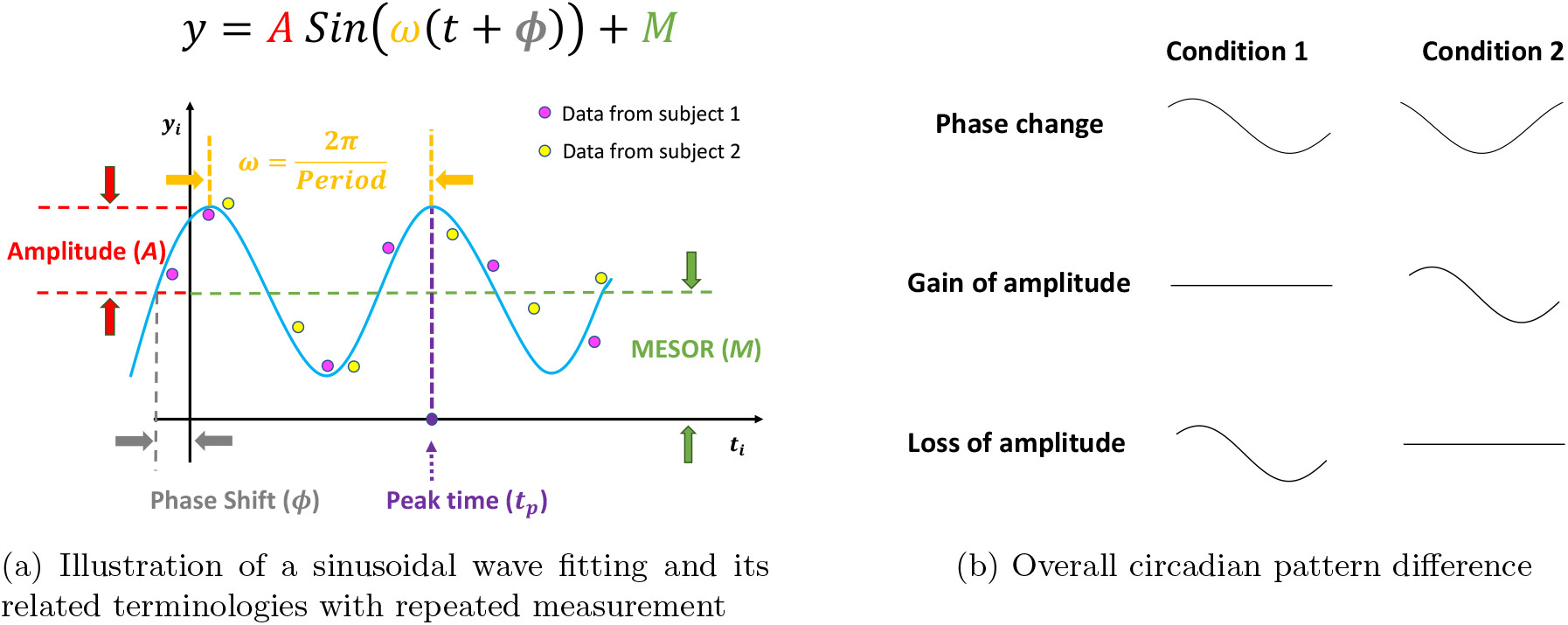
Illustration of circadian rhythmicity detection and differential circadian analysis. (a) Illustration of a sinusoidal wave fitting and its related terminologies with repeated measurement. (b) Three types of overall circadian pattern difference comparing two experimental conditions.

### 2.2 Circadian rhythmicity detection with repeated measurement

In this section, we develop likelihood-based methods to examine the existence of a circadian rhythmicity within one experimental condition. Denote ***y***_***i***_ = (*y*_*i*1_, …, *y*_*ik*_, …, *y*_*iK*_) as the expression values of a gene for subject *i*(1 ≤ *i* ≤ *n*), where *n* is the total number of subjects. The *i*^*th*^ subject is repeatedly measured *K* times, with the gene expression value at circadian time *t*_*ik*_ being *y*_*ik*_ (See purple or yellow dots in Figure 1a). For the ease of method description, we assume all subjects are measured the same *K* times. But our method can be easily extended to accommodate different number of repeated measurement for different subjects, which has been implemented in our software package. We assume:

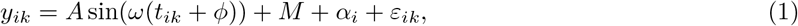

where 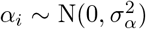 is the subject specific random effect, and 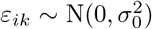 is the error term for subject *i* at circadian time *t*_*ik*_; *α*_*i*_ and *ε*_*ik*_ are independent. Under this formulation,

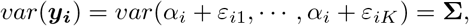

where **Σ** ∈ ℝ^*K*×*K*^ is the variance-covariance matrix, with the diagonal elements being 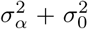 and the off-diagonal elements being 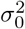. Therefore, the correlation structure from the repeated measurement of the same subject is taken into account by the off diagonal elements of **Σ**. To benchmark the goodness of fit of the circadian rhythmicity, we adopt *r* = *A/σ*_0_, which has been shown to be the key effect size parameter for circadian power calculation (Zong et al., 2022).

Based on these assumptions, we derive procedures for examining circadian rhythmicity, with the null hypothesis being there exists no circadian rhythmicity (i.e., *H*_0_ : *A* = 0) and the alternative hypothesis being there exists a circadian rhythmicity (i.e., *H*_*A*_ : *A* ≠ 0). For the ease of discussion, we re-write Equation 1 as

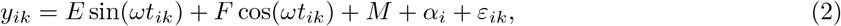

where *E* = *A* cos(*ωϕ*), and *F* = *A* sin(*ωϕ*). Then the hypothesis testing framework is equivalent to:

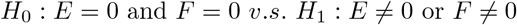

We denote this method as Rpt rhthmicity throughout this manuscript.

#### 2.2.1 Likelihood ratio test for Rpt rhthmicity

Denote *l* as the log likelihood function of Equation 2 and ***β*** = (*E, F*)^⊤^. The null hypothesis is *H*_0_ : ***β*** = ***β***_0_ = (0, 0)^⊤^, and the corresponding likelihood is 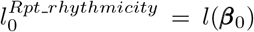. The alternative hypothesis is 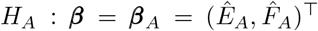, and the corresponding likelihood is 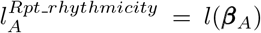, where ***β***_*A*_ is the maximum likelihood estimate of Equation 2 under *H*_*A*_. The likelihood ratio test statistic is: 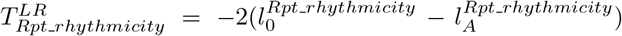. Since the degree of freedom is 2, under *H*_0_, 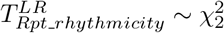.

#### 2.2.2 Kenward-Roger F test for Rpt rhthmicity

Since the likelihood ratio test is an asymptotic test, it may result in inflated type I error rate when the sample size is small (Pinheiro and Bates, 2006). We also implement the Kenward-Roger F test (Halekoh and Højsgaard, 2014) to perform statistical inferences, which has been shown to produce type I error rate that is close to its nominal level and to be less sensitive to sample size (Luke, 2017) for linear mixed model.

#### 2.2.3 Other competing methods

To benchmark the performance, we also compare our proposed likelihood-based mixed model method to some other existing methods that cannot model the repeated measurement data, including LR_rhythmicity (Ding et al., 2021), Lomb-Scargle periodograms (Glynn et al., 2006), JTK CYCLE (Hughes et al., 2010) and MetaCycle (Wu et al., 2016).

### 2.3 Differential circadian analysis with repeated measurement

In this section, we develop likelihood-based testing procedures to identify genes showing differential circadian patterns. In this paper, we will focus on examine whether there is a difference in the overall circadian pattern under the repeated measurement setting. As shown in Figure 1b, the overall circadian pattern difference is consisted of gain of rhythmicity, loss of rhythmicity, and phase change. In this paper, we don’t consider MESOR change as a differential circadian pattern, since it can also be categorized as differential expression analysis (Ritchie et al., 2015; Robinson et al., 2010; Love et al., 2014).

Denote *y*_1*ik*_ as the gene expression value of subject *i*(1 ≤ *i* ≤ *n*_1_) at circadian time *t*_1*ik*_ in experimental condition 1, where *n*_1_ is the number of subjects in condition 1, *k*(1 ≤ *k* ≤ *K*) is the *k*^*th*^ measurement for subject *i*, and *K* is the total number of repeated measurement. *y*_2*jk*_ is the gene expression value of subject *j*(1 ≤ *j* ≤ *n*_2_) at circadian time *t*_2*jk*_ in experimental condition 2, where *n*_2_ is the number of subjects in condition 2. Note that *y*_1*ik*_ and *y*_2*jk*_ are from the same gene, but under different experimental conditions. We assume the following models:

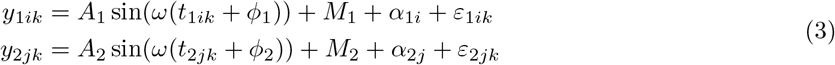

where 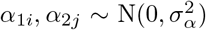 are the subject specific random effects; 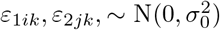 are the error terms; These terms, including *α*_1*i*_, *α*_2*j*_, *ε*_1*ik*_, and *ε*_2*jk*_, are assumed to be independently and identically distributed. As explained in Section 2.2, the correlation structure of the repeated measurements from the same subject is taken into account via the subject specific random effect. *A*_1_, *ϕ*_1_, and *M*_1_ are the amplitude, phase, MESOR for the experimental condition 1, and *A*_2_, *ϕ*_2_, and *M*_2_ are for experimental condition 2. Though our method is based on comparing two experimental conditions, our linear mixed model framework can be further extended when the experimental condition is a continuous variable. Below we state the null hypothesis and the alternative hypothesis for testing differential circadian patterns, based on Equation 3.

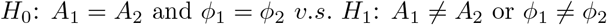

For the ease of discussion, we re-write Equation 3 as

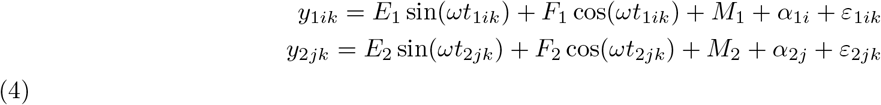

Then the hypothesis testing framework is equivalent to:

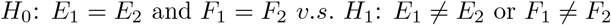

We denote this method as Rpt_diff throughout this manuscript.

#### 2.3.1 Likelihood ratio test for Rpt_diff

Denote 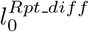 as the log likelihood for Equation 4 under the null (i.e., *E*_1_ = *E*_2_ and *F*_1_ = *F*_2_); 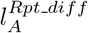 as the log likelihood under the null (i.e., *E*_1_ ≠ *E*_2_ or *F*_1_ ≠ *F*_2_). The likelihood ratio test statistic is: 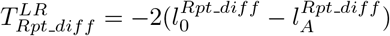. Since the degree of freedom is 2, under 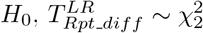.

#### 2.3.2 Kenward-Roger F test for Rpt_diff

Again, in order to address the issue of inflated type I error rate for the likelihood ratio test because of small sample sizes, we also adopt the KR F-test.

#### 2.3.3 Competing methods for differential circadian analysis

To benchmark the performance, we also compare the performance of our method with other existing methods, including RobustDODR (Thaben and Westermark, 2016), HANOVA (Thaben and Westermark, 2016) and LimoRhyde (Singer and Hughey, 2019), which are all designed to examine the difference of the overall circadian pattern between two experimental conditions. However, none of these methods can model the repeated measurement data.

## 3 Simulation

We simulated repeatedly measured gene expression data as well as their circadian time. We evaluated the performance of (i) Rpt_rhythmicity (i.e., circadian rhythmicity detection with repeated measurement) and (ii) Rpt_diff (i.e., differential circadian analysis with repeated measurement) in terms of both type I error rate and statistical power (i.e., 1 – type II error rate). To benchmark the performance, we also compared to their completing methods which could not model the repeated measurement.

### 3.1 Simulation for circadian rhythmicity detection

#### 3.1.1 Simulation settings

Denoted *i*(1 ≤ *i* ≤ *n*) as the subject index and *n* as number of subjects. Each subject *i* was repeatedly measured *K* times, and circadian time *t*_*ik*_ (1 ≤ *k* ≤ *K*) were evenly distributed within a 24-hour window (i.e. *t*_*i*1_ = 24*/K, t*_*i*2_ = 2 × 24*/K*, …, *t*_*iK*_ = 24). We simulated the gene expression value for sample *i* at circadian time *t*_*ik*_ using Equation 1. Our basic parameter setting is listed below. For each gene, we set the number of subjects *n* = 12; and the number of repeated measurements for each subject *K* = 12. The circadian time were sampled every 2 hours (i.e. *t*_*i*1_ = 2, *t*_*i*2_ = 4, …, *t*_*i*12_ = 24). Though we sampled integer circadian time and evenly spaced interval time, our methods could take non-integer circadian time with any distribution as input. For each gene *g*(1 ≤ *g* ≤ *G*), we set amplitude *A* = 1; phase *ϕ* ∼ UNIF(0, 12); MESOR *M* ∼ UNIF(0, 3); the error terms (*α*_*i*_+*ε*_*i*1_, …, *α*_*i*_+*ε*_*iK*_) were generated from multivariate normal distribution MVN(0, **Σ**), where **Σ** was the variance-covariance matrix with 1 as the diagonal elements and *ρ* = 0.2 as the off-diagonal elements. We simulated *G* = 10, 000 genes for each simulation, and each simulation was repeated *B* = 10 times to increase number of replications and to obtain an estimate of standard deviation. To examine the impact of sample size *n*, number of repeated measurements *K*, and within-subject correlation among repeated measurements *ρ*, we further simulated following variations.

1. Impact of sample size. We varied *n* = 6, 12, 24 while fixing other parameters in the basic parameter setting.
2. Impact of number of observations. We varied number of repeated measurements *K* = 12, 24, 48 for each subject, while fixing other parameters in the basic parameter setting.
3. Impact of within-subject correlation among observations. We varied **Σ**’s off-diagonal term *ρ* = 0, 0.2, 0.5 while fixing other parameters in the basic parameter setting.

#### 3.1.2 Type I error rate for Rpt_rhythmicity

We evaluated our proposed linear mixed model methods for the repeatedly measured circadian rhythmicity detection, including both the likelihood ratio test (i.e., Rpt_rhythmicity (LR)) and the F test (i.e., Rpt_rhythmicity (F)) in terms of type I error rate. To benchmark the performance, we also compared to several existing methods without considering repeated measurements, including Lomb-Scargle periodograms, JTK CYCLE, MetaCycle and LR_rhythmicity in detecting circadian rhythmicity. As shown in Figure 2, when varying (i) number of subject *n* (Figure 2a); (ii) number of repeated measurements *K* (Figure 2b), Rpt_rhythmicity (LR) and Rpt_rhythmicity (F) controlled type I errors around the nominal level (i.e. 5%), while all other competing methods had conservative type I errors. We also noted that the Rpt_rhythmicity (LR) has a slightly inflated type I error (≈ 0.055) when sample size was small (*n* = 6), and was more accurate when sample size was large. On the other hand, Rpt_rhythmicity (F) was less sensitive to the variation of sample sizes. This indicates that the F Test should be preferred when the sample size is small. When varying (iii) within-subject correlation *ρ*, in addition to Rpt_rhythmicity (LR) and Rpt_rhythmicity (F), LR_rhythmicity also successfully controlled the type I error rate to the 5% nominal level when *ρ* = 0, which was not unexpected since LR_rhythmicity was based a cosinor model without considering correlations among observations. In summary, under the repeated measurement setting, only the Rpt_rhythmicity (LR) and the Rpt_rhythmicity (F) could control the type I error to the 5% nominal level, while the other competing methods without considering repeated measurement will result in an inflated type I error rate.

**Figure 2:**
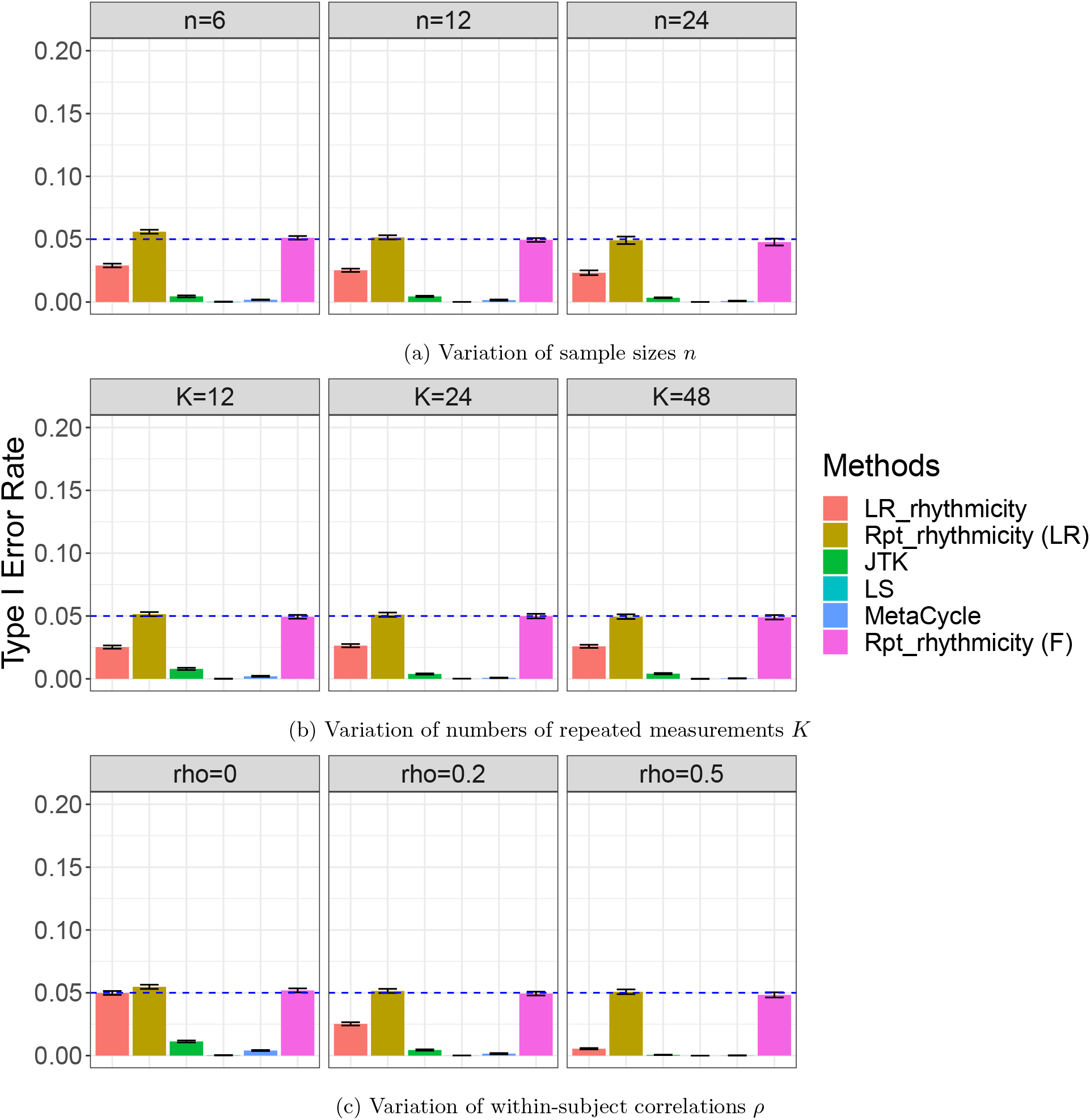
Type I error rate at nominal *α* level 5% for 6 different methods in detecting circadian rhythmicity under the repeated measurement setting. Rpt_rhythmicity (LR) is our proposed linear mixed model method using likelihood ratio test; Rpt_rhythmicity (F) is the linear mixed model with Kenward-Roger F-Test, which is less sensitive to sample sizes. In Figure 2a, the sample sizes were varied at *n*=6, 12 and 24. The blue dashed line is the 5% nominal level. A higher than 5% blue dashed line bar indicates an inflated type I error rate; a lower than 5% blue dashed line bar indicates a conservative type I error rate; and a bar at the blue dashed line indicates an accurate type I error rate (i.e., *p* = 0.05). In Figure 2b, the numbers of repeated measurements were varied at *K*=12, 24 and 48. In Figure 2c, the within-subject correlations were varied at *ρ* = 0, 0.2 and 0.5.

#### 3.1.3 Power analysis for Rpt_rhythmicity

In terms of power analysis, we only included methods that could correctly controlled type I error rate at the 5% nominal level, otherwise the power was not comparable because we would not know if the difference in power was caused by the test procedure itself or inflated/conservative type I error rate. In our case, we compared powers of Rpt_rhythmicity (LR), Rpt_rhythmicity (F) and LR_rhythmicity. Figure 3a shows the powers of the three methods with varying number of subjects *n*. Powers increased when the number of subjects increased, and Rpt_rhythmicity (LR) and Rpt_rhythmicity (F) were more powerful than LR_rhythmicity.

**Figure 3:**
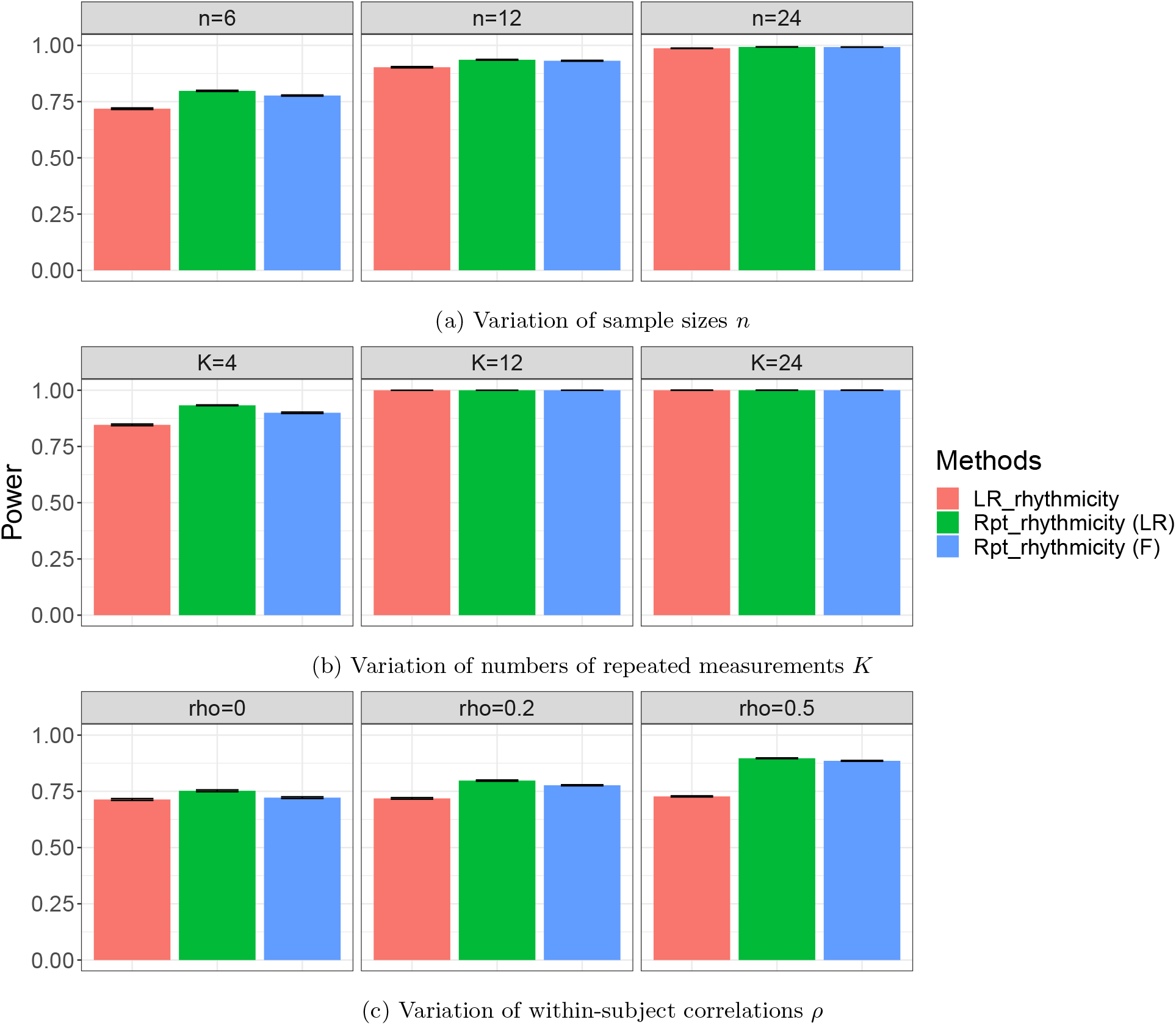
Power analysis for 3 different methods in detecting circadian rhythmicity under the repeated measurement setting. Rpt_rhythmicity (LR) is our proposed linear mixed model method using likelihood ratio test; Rpt_rhythmicity (F) is the linear mixed model with Kenward-Roger F-Test, which is less sensitive to sample sizes. In Figure 3a, the sample sizes were varied at n=6, 12 and 24. In Figure 3b, the numbers of repeated measurements were varied at K=12, 24 and 48. In Figure 3c, the within-subject correlations were varied at *ρ* = 0, 0.2 and 0.5.

Similar results were obtained when varying number of repeated measurement *K* (Figure 3b). Figure 3c showed the power for varying *ρ*. When *ρ* = 0, all three methods were equally powerful; while when *ρ* > 0, both Rpt_rhythmicity (LR) and Rpt_rhythmicity (F) were more powerful than LR_rhythmicity. These results were expected since Rpt_rhythmicity could capture the within-subject correlation structure. In summary, under the repeated measurement setting, our proposed the Rpt_rhythmicity (LR) and the Rpt_rhythmicity (F) are the most powerful methods in detecting circadian rhythmicity.

#### 3.1.4 Power analysis for Rpt_rhythmicity

In terms of power analysis, we only included methods that could correctly controlled type I error rate at the 5% nominal level, otherwise the power might not be comparable because we would not know if the difference in power was caused by the test procedure itself or inflated/deflated type I error rate. In our case, we compared powers of Rpt_rhythmicity (LR), Rpt_rhythmicity (F) and LR_rhythmicity. Figure 3a shows the powers of the three methods with varying number of subjects *n*. Powers increased when the number of subjects increased, and Rpt_rhythmicity (LR) and Rpt_rhythmicity (F) were more powerful than LR_rhythmicity. Similar results were obtained when varying number of repeated measurement *K* (Figure 3b. Figure 3c) showed the power for varying *ρ*. When *rho* = 0, all three methods were equally powerful; while when *ρ* > 0, both Rpt_rhythmicity (LR) and Rpt_rhythmicity (F) were more powerful than LR_rhythmicity. These results were expected since Rpt_rhythmicity could capture the within-subject correlation structure. In summary, under the repeated measurement setting, our proposed the Rpt_rhythmicity (LR) and the Rpt_rhythmicity (F) are the most powerful methods in detecting circadian rhythmicity.

### 3.2 Simulation for differential circadian analysis

#### 3.2.1 Simulation settings

For differential circadian analysis, the basic simulation setting was based on Equation 3, in which we set *n*_1_ = *n*_2_ = 12; *K* = 12. For each gene, amplitudes *A*_1_ = *A*_2_ = 1, phases *ϕ*_1_ = *ϕ*_2_ = 5, MESOR *M*_1_ = *M*_2_ = 3 and total error terms (random intercept + residual error) are generated from multivariate normal distribution (*α*_1*i*_ + *ε*_1*i*1_, …, *α*_1_*i* + *ε*_1*iK*_) ∼ MVN(0, **Σ**_**1**_) and (*α*_2*i*_ + *ε*_2*i*1_, …, *α*_2*i*_ + *ε*_2*iK*_) ∼ MVN(0, **Σ**_**2**_), where **Σ**_**1**_ = **Σ**_**2**_ were the variance-covariance matrices with 1 as the diagonal elements and *ρ* = 0.2 as the off-diagonal elements. Again, we simulated *G* = 10, 000 genes for each simulation, and each simulation was repeated *B* = 10 times to increase number of replications and to obtain an estimate of standard deviation. We further simulated following variations to examine the impact of number of subject *n*, number of repeated measurement *K*, and within-subject correlation *ρ*.

1. Impact of number of subjects. We varied *n* = 6, 12, 24 in both experiment conditions while fixing other parameters in the basic parameter setting.
2. Impact of number of repeated measurement for each subject. We varied number of repeated measurements *K* = 12, 24, 48 in both experiment conditions for each subject, while fixing other parameters in the basic parameter setting.
3. Impact of within-subject correlation among observations. We varied **Σ**_**1**_ and **Σ**_**2**_’s off-diagonal term *ρ* = 0, 0.2, 0.5 while fixing other parameters in the basic parameter setting.

#### 3.2.2 Type I error rate for Rpt_diff

We evaluated our proposed linear mixed model methods for repeatedly measured differential circadian analysis, including both the likelihood ratio test (i.e., Rpt_diff (LR)) and the F test (i.e., Rpt_diff (F)) in terms of type I error rate. To benchmark the performance, we also compared to several existing methods for differential circadian analysis without considering repeated measurements, including HANOVA, RobustDODR and Limorhyde. Figure S1a showed the variation of type I error rates with respect to different number of subjects, where all methods controlled type I errors around nominal level (i.e. 5%). We also observed that the Rpt_diff (LR) had a slightly inflated type I error rate when the number of subjects was small (e.g., *n* = 6), while Rpt_diff (F) was less sensitive to sample size. Similarly, all methods could control type I errors around 5% nominal level when varying number of repeated measurement *K* (Figure S1b) or varying within-subject correlation among repeated measurements *ρ* (Figure S1c). We noticed that, in Figure S1c, HANOVA and RobustDODR had slightly conservative type I error rates when within-subject correlation increased. To summarize, all these methods can control the type I error around the 5% nominal level under the repeated measurement simulation setting.

#### 3.2.3 Power analysis for Rpt_diff

In terms of power analysis, we included all pre-mentioned methods since they all controlled the type I errors around the 5% nominal level. As shown in Figure 4a, powers increased for all methods when number of subject *n* increased. In addition, our proposed method Rpt_diff (LR) and Rpt_diff (F) were more powerful compared to their competing methods. Similar results were obtained by varying *K* – number of repeated measurement for each subject (Figure 4b). When varying the within-subject correlation *ρ* (Figure 4c), we observe that if *ρ* = 0, Rpt_diff (LR), Rpt_diff (F) and Limorhyde were the most powerful methods; if *ρ* > 0, Rpt_diff (LR) and Rpt_diff (F) were more powerful, which was expected since only these two methods took into account the repeated measurement. In summary, under the repeated measurement setting, our proposed the Rpt_diff (LR) and the Rpt_diff (F) are the most powerful methods in detecting differential circadian patterns.

**Figure 4:**
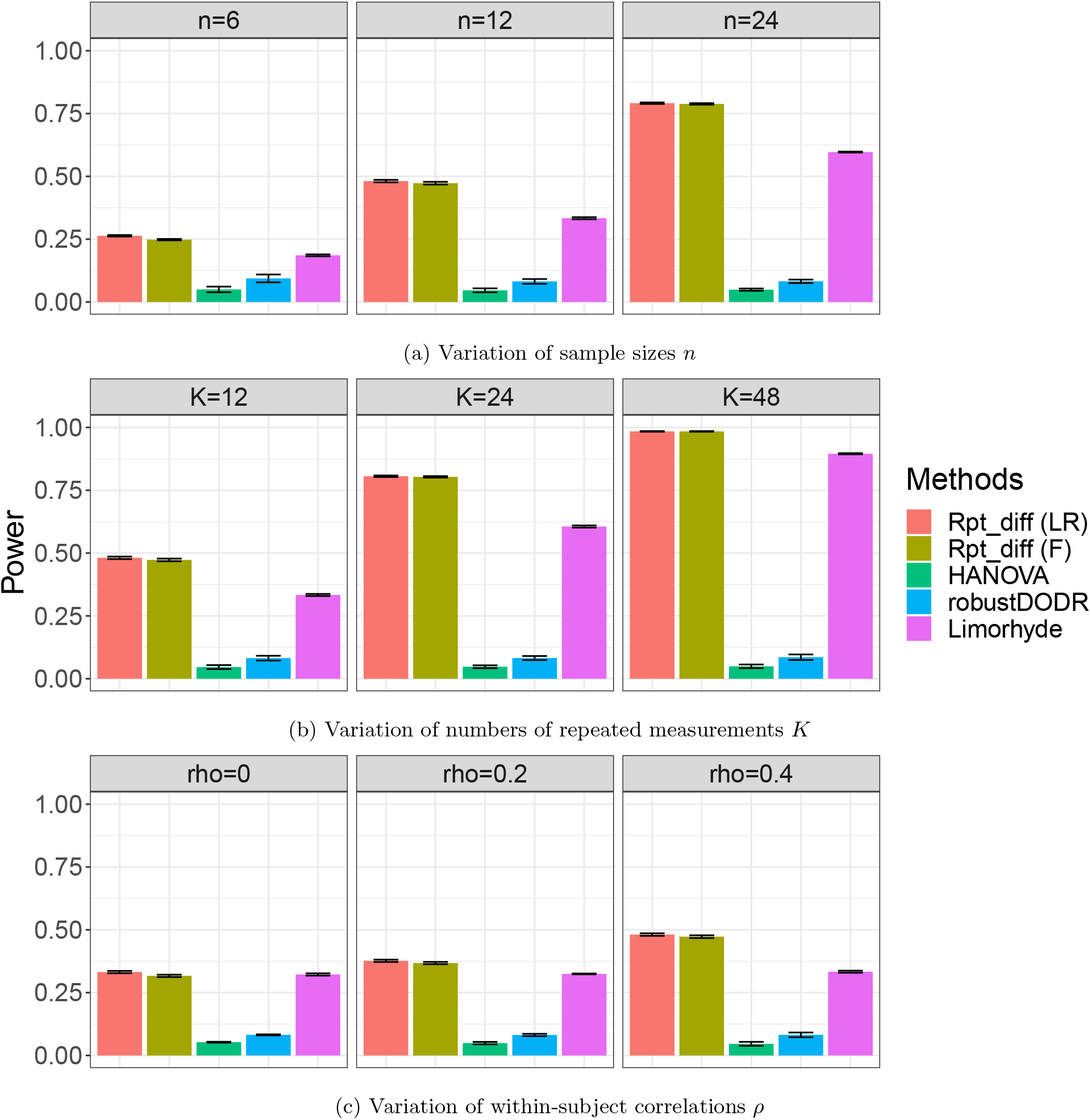
Power analysis for 5 different methods in detecting differential circadian pattern with repeated measurement. Rpt_diff (LR) represents our proposed linear mixed model method using likelihood ratio test. Rpt_diff (F) indicates the linear mixed model with Kenward-Roger F-Test, which is less sensitive to sample sizes. In Figure 4a, the sample sizes were varied at n=6, 12 and 24. In Figure 4b, the numbers of repeated measurements were varied at K=12, 24 and 48. In Figure 4c, the within-subject correlations were varied at *ρ* = 0, 0.2 and 0.4.

## 4 Real data applications

In this section, we applied our repeated measurement methods (Rpt_rhythmicity and Rpt_diff) in two real circadian transcriptomic data applications, including (i) a human restricted feeding study via RNA Sequencing data comparing restricted feeding conditions versus unrestricted feeding conditions; and (ii) a human sleep restriction study via microarray data comparing sufficient sleep conditions versus insufficient sleep conditions. To benchmark the performance, we also compared our proposed methods to other competing methods (without considering repeated measurement). Let *p*_*r*_ denote the p-value from circadian rhythmicity detection (i.e. from Rpt_rhythmicity), we used *p*_*r*_ < 0.01 as statistical significance cutoff to declare circadian rhythmicity unless otherwise specified. Since the pre-requisite for the differential circadian pattern detection is that there exists circadian rhythmicity in at least one experimental condition, we recommend perform a filtering step prior to the differential circadian analysis, and only keep the candidate genes with *p*_*r*_ < 0.01 in at least one experimental condition. Let *p*_*d*_ denote the p-value from differential circadian pattern detection (i.e. from Rpt_diff), and we used *p*_*d*_ < 0.01 as statistical significance cutoff to declare differential circadian pattern unless otherwise specified.

### 4.1 Human time-restricted feeding data

We evaluated the performance of our proposed repeated measurement methods in an RNA Sequencing data of human skeletal muscle tissue. Detailed description of this study has been described elsewhere (Lundell et al., 2020). To be brief, this study included 11 men (30 - 45 years old) with overweight or obesity. A crossover design was employed, where all participants were randomized into a time-restricted feeding (TRF) group or an unrestricted feeding (URF) group in the first period. After a wash-out period, all participants were assigned to the other group they haven’t gone through in the second period. For each period, the skeletal muscle samples of each participant were repeatedly measured every 4 hours within a 24 hours period. The time of sample collection was used as the circadian time. Although there were some missing values, each participant was repeatedly measured 4-6 times in each period, resulting in a total of 63 samples in TRF group and 62 samples in URF group. The dataset is publicly available on GEO (GSE129843). The gene expression count data were normalized using “count per million reads” (cpm). After filtering out genes with mean cpm less than 1, 13,167 genes remained for the analysis. We further performed log2 transformation (i.e. log2(x+1), x is the cpm of a gene) to improve normality of the data. Since the sample size is large, we employed the likelihood ratio test for statistical inference.

#### 4.1.1 Circadian rhythmicity detection

To detect genes showing circadian rhythmicity, we applied our proposed Rpt_rhythmicity to this time restricted feeding data. With *p*_*r*_ < 0.01 cutoff, we identified 2,078 and 1,605 circadian genes for the TRF group and the URF group, respectively. Figure 5a and 5b showed 6 core circadian genes, including *ARNTL, DBP, NR1D1, PER1, PER2* and *PER3* in the TRF group and the URF group. These 6 circadian genes are known to be clock-regulated (Zhang et al., 2014), and the Rpt_rhythmicity reported highly significant p-values (1.51 × 10^−22^ ∼ 4.69 × 10^−05^). To benchmark the performance, we also applied the LR_rhythmicity (Ding et al., 2021), which is based on the cosinor model without considering the repeated measurement. It identified 1,407 and 935 circadian genes for the TRF group and the URF group, respectively, under the same *p*_*r*_ < 0.01 cutoff. In addition, the p-values of the 6 core circadian genes using the LR_rhythmicity became less significant (1.65 × 10^−21^ ∼ 7.99 × 10^−05^). The less numbers and levels of significant circadian genes were expected, since our proposed Rpt_rhythmicity is more powerful under the repeated measurement setting.

**Figure 5:**
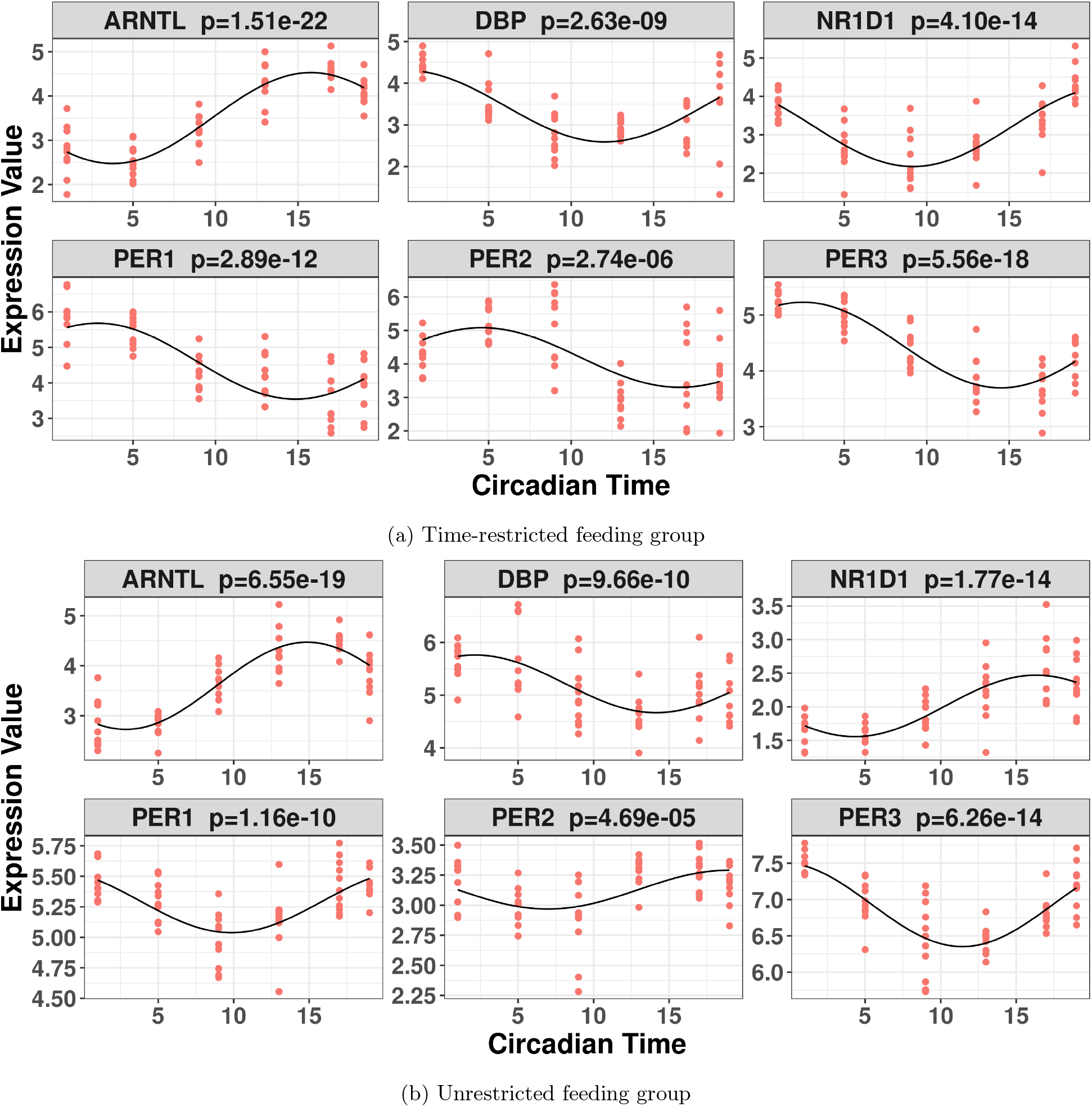
Visualization of the 6 core circadian genes in the human restricted feeding data. The p-values were obtained using the Rpt_rhythmicity.

#### 4.1.2 Differential circadian pattern detection

We applied the Rpt_diff to perform differential circadian analysis comparing the TRF to the URF group. Before applying the Rpt_diff, we first used the Rpt_rhythmicity and obtained 2,850 candidate genes showing circadian rhythmicity (*p*_*r*_ < 0.01) in either the TRF group or the URF group. Then with *p*_*d*_ < 0.01 cutoff, we identified 230 genes showing differential patterns. Figure 6 showed top four most significant differential circadian genes. For example, gene *KLHL25* (*p*_*d*_ = 5.61 × 10^−09^) in Figure 6a showed reduced amplitude and shifted phase in the URF group compared to the TRF group. To benchmark the performance, we also compared to other differential circadian methods without considering repeated measurement, including HANOVA, RobustDODR and LimoRhyde. Based on the same set of 2,850 candidate genes and under *p*_*d*_ < 0.01 cutoff, the HANOVA, RobustDODR and LimoRhyde identified 94, 103 and 96 significant differential circadian genes respectively, which were much less than the 230 genes detected by the Rpt_diff. This is expected because the Rpt_diff can take advantage of the within subject correlation from the repeated measurement data to improve statistical power.

**Figure 6:**
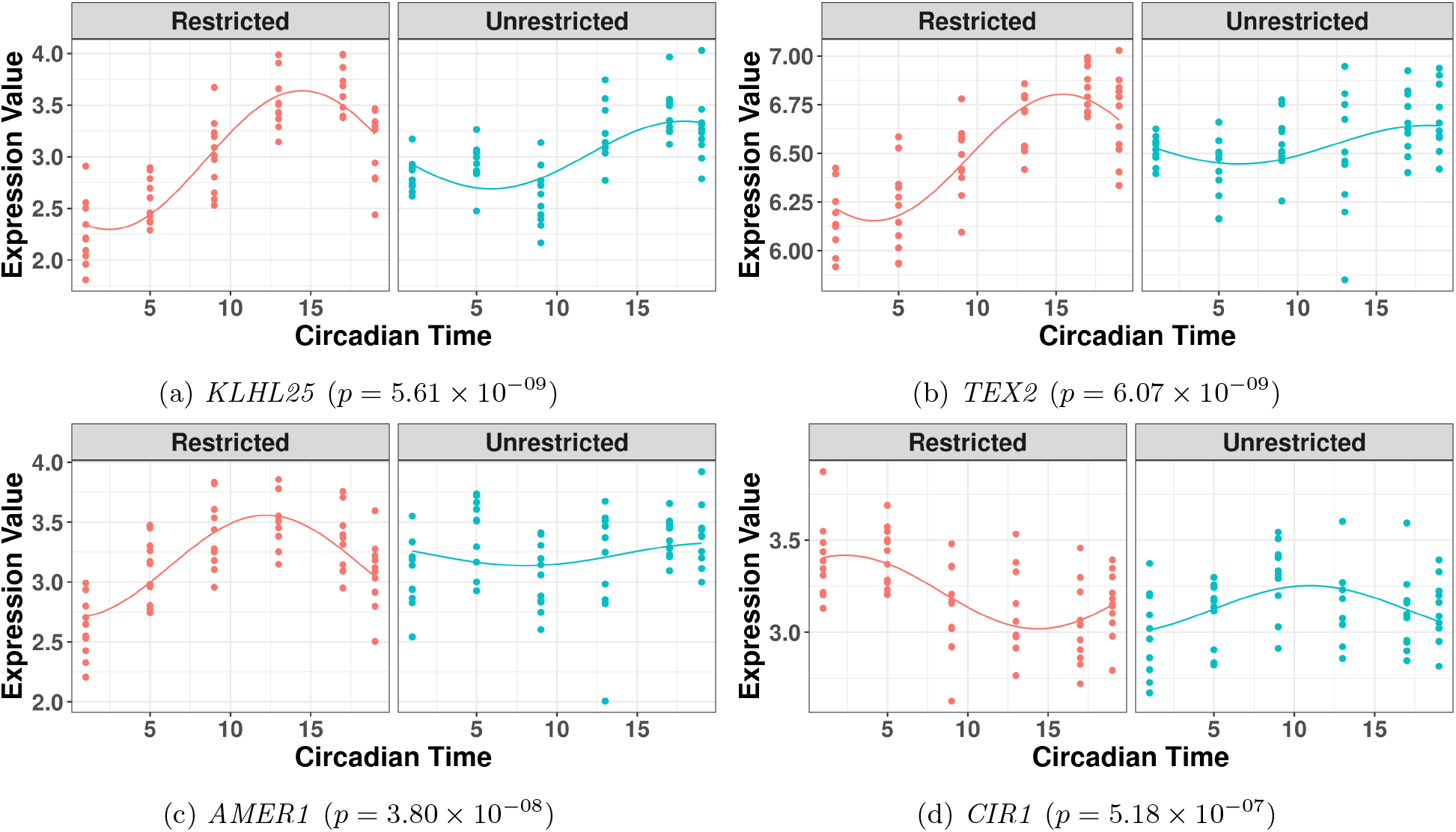
Top four significant genes showing differential circadian patterns from the human time-restricted feeding data. The two experimental conditions are the time-restricted feeding (TRF) group and the unrestricted feeding (URF) group

### 4.2 Human sleep restriction data

We further evaluated the performance of our proposed repeated measurement methods in a gene expression microarray data of human blood. In this study, a total of 438 samples (217 with insufficient sleep and 221 with sufficient sleep) were collected from 26 participants (Möller-Levet et al., 2013). In this cross over design study, each participant was assigned to a week intervention of both sufficient sleep (> 6 hours per day) and insufficient sleep (6 hours per day) with a random order, separated by a 10-days interval. Following each of these one-week interventions, all participants were kept awake for a day, a night and the subsequent day. 10 blood samples were taken while subjects were in each of these periods of extended wakefulness. The time of sample collection was used as the circadian time. The dataset is publicly available on GEO (GSE39445). In this microarray data, if multiple probes match to the same gene symbol, we will use their median value to represent the gene expression level. After this step, 19,541 gene were included in the analysis. Since the sample size is large, we employed the likelihood ratio test for statistical inference.

#### 4.2.1 Circadian rhythmicity detection

We applied the Rpt_rhythmicity to this human sleep restriction data with repeated measurement to detect circadian rhythmicity. Since the total sample size in this study is fairly large, we also adopted the circadian effect size parameter *r* = *A/σ*_0_ (Zong et al., 2022) in addition to the *p*_*r*_ criteria to declare genes showing circadian rhythmicity. Under effect size *r* < 0.4 and *p*_*r*_ < 0.01 criteria, we identified 2,743 and 2,851 circadian genes under the sufficient sleep condition and the insufficient sleep condition, respectively. In terms of the 6 core circadian genes (i.e., *ARNTL, DBP, NR1D1, PER1, PER2* and *PER3*), our method (Rpt_rhythmicity) reported highly significant p-values (3.56 × 10^−24^ ∼ 2.72 × 10^−02^), indicating strong ability of circadian rhythmicity detection. These 6 genes are shown in Figure S2a (for the sufficient sleep condition) and Figure S2b (for the insufficient sleep condition). To benchmark the performance, we also applied the LR_rhythmicity without considering the repeated measurement. Under the same significance criteria, the LR_rhythmicity identified 30 and 46 genes for the sufficient sleep condition and the insufficient sleep condition, respectively. In addition, the significance level of the 6 core circadian genes became less significant, with p-values ranging from 2.18 × 10^−11^ to 2.95 × 10^−01^. Collectively, these results indicate the Rpt_rhythmicity is more powerful in detecting circadian rhythmicity under the repeated measurement setting than the non-repeated measurement methods.

#### 4.2.2 Differential circadian pattern detection

We further applied the Rpt_diff to perform differential circadian analysis comparing the sufficient sleep condition and the insufficient sleep condition. Before applying the Rpt_diff, we first selected candidate genes showing circadian rhythmicity in either the sufficient sleep condition or the insufficient sleep condition (n = 4,602) from the previous step. Then with *p*_*d*_ < 0.01 cutoff, we identified 61 genes showing differential pattern. Figure S3 showed top four most significant genes showing differential circadian pattern. For example, gene *PSTPIP1* in the sufficient sleep condition showed increased amplitude and shifted phase compared to the insufficient sleep condition (*p*_*d*_ = 3.09 × 10^−04^, Figure S3a). To benchmark the performance, we further compare with other differential circadian methods. Under *p*_*d*_ < 0.01 cutoff, the HANOVA, RobustDODR and LimoRhyde only identified 4, 1 and 4 significant differential circadian genes respectively, which were much less than the 61 genes detected by the Rpt_diff. This is not surprising since these competing methods cannot take into account the within subject correlation from the repeated measurement data.

## 5 Discussion

In summary, we developed linear mixed model based methods for detecting (i) circadian rhythmicity (i.e., Rpt_rhythmicity) and (ii) differential circadian pattern (i.e., Rpt_diff) for repeatedly measured circadian data. The within subject correlation from the repeated measurement data is modeled by the subject-specific random effect. We developed likelihood ratio test and F test as statistical inference methods to obtain p-values. To our knowledge, our methods are the first to enable modeling repeatedly measured data in the circadian field, which will make a huge impact as such data are becoming increasingly prevalent in the circadian field. Simulations showed that both our methods (Rpt_rhythmicity and Rpt_diff) could successfully control type I error at 5% nominal level (i.e. produces an accurate p-value) and have higher power than other competing methods when the circadian data were repeated measured. We also applied our methods in repeatedly measured transcriptomic data applications including a human time restricted feeding data and a human sleep restriction data. Superior performance of our methods has been observed in these applications compared to their competing methods.

Our methods have the following strengths. (i) Our methods are the first to enable modeling the repeated measurement design in circadian analysis. (ii) When the circadian data are repeatedly measured, our methods can not only control the type I error rate around the nominal level, but also be more powerful than competing methods that cannot model the repeated measurement. (iii) The likelihood ratio test is accurate when the sample size is large. To address the potential sample size issue, we also developed a F test method, which is less sensitive to sample size. (iv) Our methods have been implemented in R package ‘RepeatedCircadian’, which is publicly available on GitHub (https://github.com/RepeatedCircadian/RepeatedCircadian).

Our methods have the following limitations. (i) Our methods assume a cosinor model and residuals are normally distributed. Though this model is easy to interpret and enjoys accurate statistical inferences, it cannot be used for modeling irregular curve shape. Extensions of the repeated measurement framework toward complicated parametric models or non-parametric models are warranted. (ii) Though our model can accomendate repeated measurement design, the current model only allows comparing two experimental conditions. In modern epidemiology or animal study, the design can be more complicated. The phenotype type of interest could be a continuous variable or categorical variable, and other biological factors (e.g., age, sex, etc) could have a confounding impact on the circadian pattern. Future works to accommodate these complicated study designs needed. Nonetheless, this is the first model to achieve circadian modeling with repeated measurement, which may lay down the foundation to overcome these limitations and motivate future extensions.

## 6 Acknowledgement

H.D., L.M., K.E., Z.H. are supported by National Institution of Health (R01HL153042). K.E., Z.H. are supported by National Institution of Health (R01AR079220). H.D., C.X., Z.H. are supported by FL DEPT OF HLTH BIOMED RES PGM/JE KING (21K11) and Lung Cancer Research Foundation (AGR DTD 11-23-2020).

## References

Badia, P., Myers, B., Boecker, M., Culpepper, J., and Harsh, J. (1991). Bright light effects on body temperature, alertness, eeg and behavior. Physiology & behavior, 50(3):583–588.

Ballesta, A., Innominato, P. F., Dallmann, R., Rand, D. A., and Lévi, F. A. (2017). Systems Chronotherapeutics. Pharmacological Reviews, 69(2):161–199.

Blume, C., Lechinger, J., Santhi, N., Giudice, R. d., Gnjezda, M.-T., Pichler, G., Scarpatetti, M., Donis, J., Michitsch, G., and Schabus, M. (2017). Significance of circadian rhythms in severely brain-injured patients. Neurology, 88(20):1933–1941.

Cagnacci, A., Elliott, J., and Yen, S. (1992). Melatonin: a major regulator of the circadian rhythm of core temperature in humans. The Journal of Clinical Endocrinology & Metabolism, 75(2):447–452.

Chen, C.-Y., Logan, R. W., Ma, T., Lewis, D. A., Tseng, G. C., Sibille, E., and McClung, C. A. (2016). Effects of aging on circadian patterns of gene expression in the human prefrontal cortex. Proceedings of the National Academy of Sciences, 113(1):206–211.

Cornelissen, G. (2014). Cosinor-based rhythmometry. Theoretical Biology and Medical Modelling, 11(1):16.

Dallmann, R., Viola, A. U., Tarokh, L., Cajochen, C., and Brown, S. A. (2012). The human circadian metabolome. Proceedings of the National Academy of Sciences, 109(7):2625–2629.

Dijk, D.-J., Duffy, J. F., and Czeisler, C. A. (1992). Circadian and sleep/wake dependent aspects of subjective alertness and cognitive performance. Journal of sleep research, 1(2):112–117.

Ding, H., Meng, L., Liu, A. C., Gumz, M. L., Bryant, A. J., Mcclung, C. A., Tseng, G. C., Esser, K. A., and Huo, Z. (2021). Likelihood-based tests for detecting circadian rhythmicity and differential circadian patterns in transcriptomic applications. Briefings in Bioinformatics, 22(6). bbab224.

Glynn, E. F., Chen, J., and Mushegian, A. R. (2006). Detecting periodic patterns in unevenly spaced gene expression time series using lomb–scargle periodograms. Bioinformatics, 22(3):310–316.

Halekoh, U. and Højsgaard, S. (2014). A kenward-roger approximation and parametric bootstrap methods for tests in linear mixed models – the r package pbkrtest. Journal of Statistical Software, 59(9):1–32.

Hastings, M. H., Reddy, A. B., and Maywood, E. S. (2003). A clockwork web: circadian timing in brain and periphery, in health and disease. Nature Reviews Neuroscience, 4(8):649–661.

Hodge, B. A., Wen, Y., Riley, L. A., Zhang, X., England, J. H., Harfmann, B. D., Schroder, E. A., and Esser, K. A. (2015). The endogenous molecular clock orchestrates the temporal separation of substrate metabolism in skeletal muscle. Skeletal muscle, 5(1):17.

Hsu, P. Y. and Harmer, S. L. (2012). Circadian phase has profound effects on differential expression analysis. PloS one, 7(11):e49853.

Hughes, M. E., Abruzzi, K. C., Allada, R., Anafi, R., Arpat, A. B., Asher, G., Baldi, P., De Bekker, C., Bell-Pedersen, D., Blau, J., et al. (2017). Guidelines for genome-scale analysis of biological rhythms. Journal of biological rhythms, 32(5):380–393.

Hughes, M. E., DiTacchio, L., Hayes, K. R., Vollmers, C., Pulivarthy, S., Baggs, J. E., Panda, S., and Hogenesch, J. B. (2009). Harmonics of circadian gene transcription in mammals. PLoS Genet, 5(4):e1000442.

Hughes, M. E., Hogenesch, J. B., and Kornacker, K. (2010). Jtk cycle: an efficient nonparametric algorithm for detecting rhythmic components in genome-scale data sets. Journal of biological rhythms, 25(5):372–380.

Hughey, J. J. and Butte, A. J. (2016). Differential phasing between circadian clocks in the brain and peripheral organs in humans. Journal of biological rhythms, 31(6):588–597.

Jung, C. M., Khalsa, S. B. S., Scheer, F. A., Cajochen, C., Lockley, S. W., Czeisler, C. A., and Wright Jr, K. P. (2010). Acute effects of bright light exposure on cortisol levels. Journal of biological rhythms, 25(3):208–216.

Koike, N., Yoo, S.-H., Huang, H.-C., Kumar, V., Lee, C., Kim, T.-K., and Takahashi, J. S. (2012). Transcriptional architecture and chromatin landscape of the core circadian clock in mammals. Science, 338(6105):349–354.

Laloum, D. and Robinson-Rechavi, M. (2020). Methods detecting rhythmic gene expression are biologically relevant only for strong signal. PLoS computational biology, 16(3):e1007666.

Li, J. Z., Bunney, B. G., Meng, F., Hagenauer, M. H., Walsh, D. M., Vawter, M. P., Evans, S. J., Choudary, P. V., Cartagena, P., Barchas, J. D., et al. (2013). Circadian patterns of gene expression in the human brain and disruption in major depressive disorder. Proceedings of the National Academy of Sciences, 110(24):9950–9955.

Lim, A. S., Klein, H.-U., Yu, L., Chibnik, L. B., Ali, S., Xu, J., Bennett, D. A., and De Jager, P. L. (2017). Diurnal and seasonal molecular rhythms in human neocortex and their relation to alzheimer’s disease. Nature communications, 8(1):1–16.

Lim, A. S. P., Srivastava, G. P., Yu, L., Chibnik, L. B., Xu, J., Buchman, A. S., Schneider, J. A., Myers, A. J., Bennett, D. A., and De Jager, P. L. (2014). 24-Hour Rhythms of DNA Methylation and Their Relation with Rhythms of RNA Expression in the Human Dorsolateral Prefrontal Cortex. PLoS Genetics, 10(11):e1004792.

Love, M. I., Huber, W., and Anders, S. (2014). Moderated estimation of fold change and dispersion for RNA-seq data with DESeq2. Genome Biology, 15(12):550.

Luke, S. G. (2017). Evaluating significance in linear mixed-effects models in R. Behavior Research Methods, 49(4):1494–1502.

Lundell, L. S., Parr, E. B., Devlin, B. L., Ingerslev, L. R., Altıntaş, A., Sato, S., Sassone-Corsi, P., Barrès, R., Zierath, J. R., and Hawley, J. A. (2020). Time-restricted feeding alters lipid and amino acid metabolite_rhythmicity without perturbing clock gene expression. Nature Communications, 11(1):4643.

Mei, W., Jiang, Z., Chen, Y., Chen, L., Sancar, A., and Jiang, Y. (2020). Genome-wide circadian rhythm detection methods: systematic evaluations and practical guidelines. bioRxiv.

Möller-Levet, C. S., Archer, S. N., Bucca, G., Laing, E. E., Slak, A., Kabiljo, R., Lo, J. C., Santhi, N., von Schantz, M., Smith, C. P., et al. (2013). Effects of insufficient sleep on circadian rhythmicity and expression amplitude of the human blood transcriptome. Proceedings of the National Academy of Sciences, 110(12):E1132–E1141.

Mure, L. S., Le, H. D., Benegiamo, G., Chang, M. W., Rios, L., Jillani, N., Ngotho, M., Kariuki, T., Dkhissi-Benyahya, O., Cooper, H. M., and Panda, S. (2018). Diurnal transcriptome atlas of a primate across major neural and peripheral tissues. Science, 359(6381):eaao0318.

Parsons, R., Parsons, R., Garner, N., Oster, H., and Rawashdeh, O. (2020). Circacompare: a method to estimate and statistically support differences in mesor, amplitude and phase, between circadian rhythms. Bioinformatics, 36(4):1208–1212.

Perrin, L., Loizides-Mangold, U., Chanon, S., Gobet, C., Hulo, N., Isenegger, L., Weger, B. D., Migliavacca, E., Charpagne, A., Betts, J. A., Walhin, J.-P., Templeman, I., Stokes, K., Thompson, D., Tsintzas, K., Robert, M., Howald, C., Riezman, H., Feige, J. N., Karagounis, L. G., Johnston, J. D., Dermitzakis, E. T., Gachon, F., Lefai, E., and Dibner, C. (2018). Transcriptomic analyses reveal rhythmic and CLOCK-driven pathways in human skeletal muscle. eLife, 7:e34114.

Pinheiro, J. and Bates, D. (2006). Mixed-effects models in S and S-PLUS. Springer science & business media.

Ritchie, M. E., Phipson, B., Wu, D., Hu, Y., Law, C. W., Shi, W., and Smyth, G. K. (2015). limma powers differential expression analyses for RNA-sequencing and microarray studies. Nucleic Acids Research, 43(7):e47–e47.

Robinson, M. D., McCarthy, D. J., and Smyth, G. K. (2010). edgeR: a Bioconductor package for differential expression analysis of digital gene expression data. Bioinformatics, 26(1):139–140.

Ruben, M. D., Wu, G., Smith, D. F., Schmidt, R. E., Francey, L. J., Lee, Y. Y., Anafi, R. C., and Hogenesch, J. B. (2018). A database of tissue-specific rhythmically expressed human genes has potential applications in circadian medicine. Science Translational Medicine, 10(458).

Sancar, A. and Van Gelder, R. N. (2021). Clocks, cancer, and chronochemotherapy. Science, 371(6524):eabb0738.

Saner, N. J., Lee, M. J.-C., Kuang, J., Pitchford, N. W., Roach, G. D., Garnham, A., Genders, A. J., Stokes, T., Schroder, E. A., Huo, Z., Esser, K. A., Phillips, S. M., Bishop, D. J., and Bartlett, J. D. (2021). Exercise mitigates sleep-loss-induced changes in glucose tolerance, mitochondrial function, sarcoplasmic protein synthesis, and diurnal rhythms. Molecular Metabolism, 43:101110.

Seney, M. L., Cahill, K., Enwright, J. F., Logan, R. W., Huo, Z., Zong, W., Tseng, G., and McClung, C. A. (2019). Diurnal rhythms in gene expression in the prefrontal cortex in schizophrenia. Nature communications, 10(1):1–11.

Singer, J. M. and Hughey, J. J. (2019). Limorhyde: a flexible approach for differential analysis of rhythmic transcriptome data. Journal of biological rhythms, 34(1):5–18.

Stenvers, D. J., Jongejan, A., Atiqi, S., Vreijling, J. P., Limonard, E. J., Endert, E., Baas, F., Moerland, P. D., Fliers, E., Kalsbeek, A., et al. (2019). Diurnal rhythms in the white adipose tissue transcriptome are disturbed in obese individuals with type 2 diabetes compared with lean control individuals. Diabetologia, 62(4):704–716.

Straume, M. (2004). Dna microarray time series analysis: automated statistical assessment of circadian rhythms in gene expression patterning. In Methods in enzymology, volume 383, pages 149–166. Elsevier.

Takahashi, J. S. (2017). Transcriptional architecture of the mammalian circadian clock. Nature Reviews Genetics, 18(3):164–179.

Thaben, P. F. and Westermark, P. O. (2014). Detecting rhythms in time series with rain. Journal of biological rhythms, 29(6):391–400.

Thaben, P. F. and Westermark, P. O. (2016). Differential rhythmicity: detecting altered rhythmicity in biological data. Bioinformatics, 32(18):2800–2808.

Wang, Y., Song, L., Liu, M., Ge, R., Zhou, Q., Liu, W., Li, R., Qie, J., Zhen, B., Wang, Y., He, F., Qin, J., and Ding, C. (2018). A proteomics landscape of circadian clock in mouse liver. Nature Communications, 9(1):1553.

Wu, G., Anafi, R. C., Hughes, M. E., Kornacker, K., and Hogenesch, J. B. (2016). Metacycle: an integrated r package to evaluate periodicity in large scale data. Bioinformatics, 32(21):3351–3353.

Yang, R. and Su, Z. (2010). Analyzing circadian expression data by harmonic regression based on autoregressive spectral estimation. Bioinformatics, 26(12):i168–i174.

Zhang, R., Lahens, N. F., Ballance, H. I., Hughes, M. E., and Hogenesch, J. B. (2014). A circadian gene expression atlas in mammals: implications for biology and medicine. Proceedings of the National Academy of Sciences, 111(45):16219–16224.

Zong, W., Seney, M. L., Ketchesin, K. D., Gorczyca, M. T., Liu, A. C., Esser, K. A., Tseng, G. C., McClung, C. A., and Huo, Z. (2022). Experimental design and power calculation in omics circadian rhythmicity detection. bioRxiv.

