## Supplementary for "Statistical Methods for Detecting Circadian Rhythmicity and Differential Circadian Patterns with Repeated Measurement in Transcriptomic Applications"

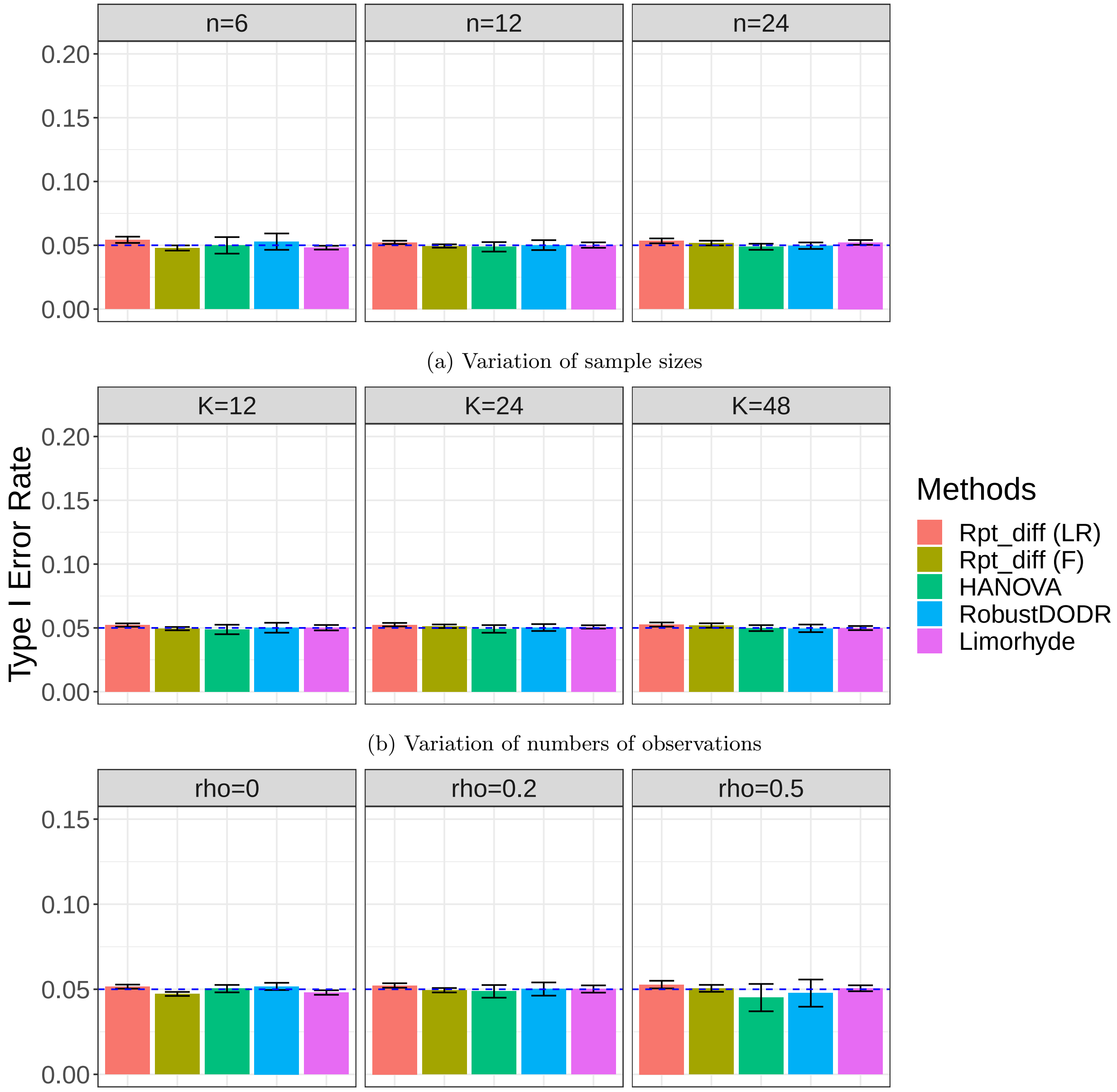


(a) Variation of sample sizes

(b) Variation of numbers of observations

(c) Variation of within-subject correlations

Figure S1: Type I error rate at nominal $\alpha$ level 5% for 5 different methods in detecting differential circadian pattern with repeated measurement. Rpt_rhythmicity (LR) represents our proposed likelihood-based method. Rpt_rhythmicity (F) indicates the linear mixed model with Kenward-Roger F-Test, which is less sensitive to sample sizes. The sample sizes were varied at n=6, 12 and 24. The numbers of observations were varied at K=12, 24 and 48. The within-subject correlations were varied at $\rho = 0$, 0.2 and 0.5. The blue dashed line is the 5% nominal level. A higher than 5% blue dashed line bar indicates an inflated type I error rate; a lower than 5% blue dashed line bar indicates a conservative type I error rate; and a bar at the blue dashed line indicates a accurate type I error rate (i.e., $p = 0.05$).

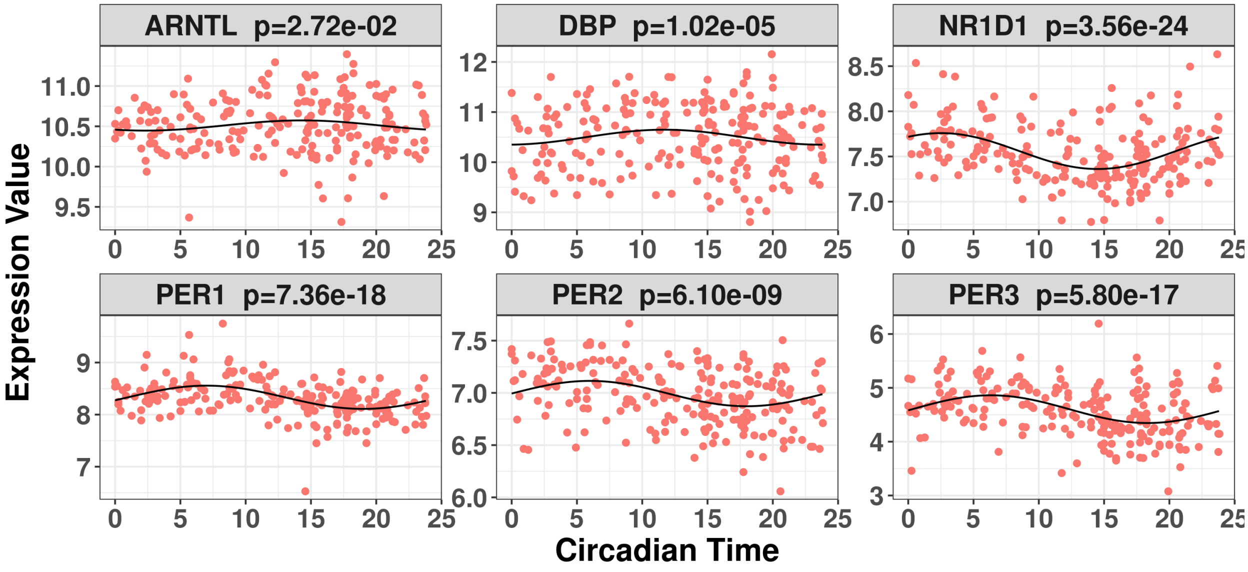


(a) Sufficient sleep condition

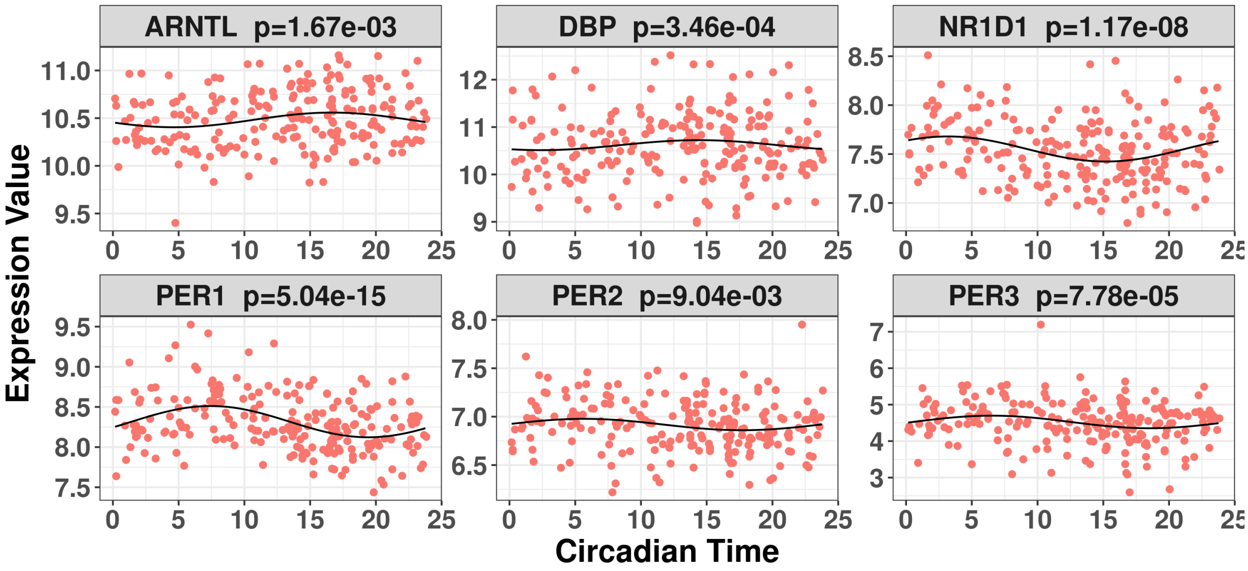


(b) Insufficient sleep condition

Figure S2: Visualization of the 6 core circadian genes in the human sufficient restriction data. The p-values were obtained using the Rpt_rhythmicity.

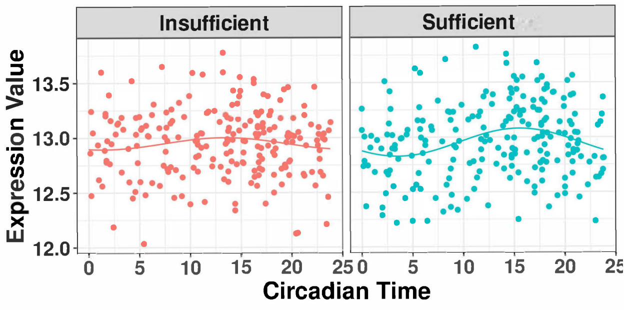


(a) *PSTPIP1* ($p = 3.09 \times 10^{-04}$)

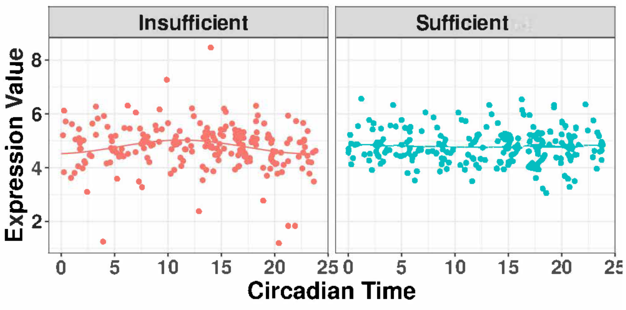


(b) *OOEP* ($p = 4.17 \times 10^{-04}$)

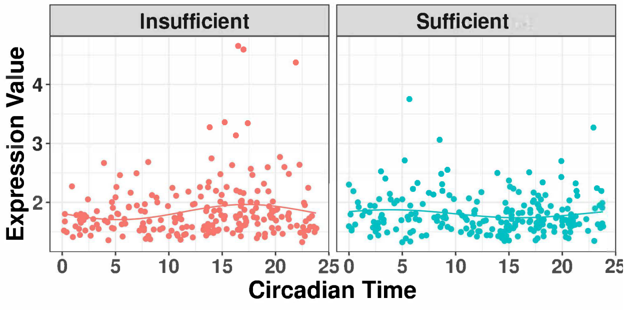


(c) *SLITRK2* ($p = 6.78 \times 10^{-04}$)

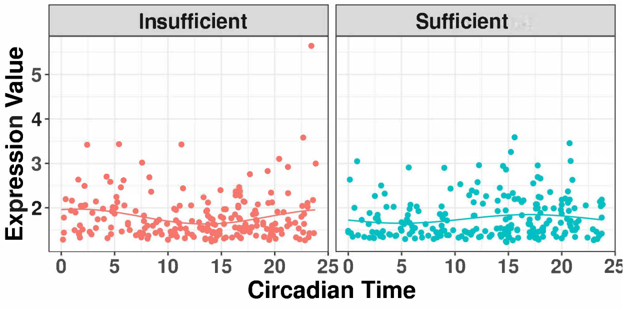


(d) *RAD21L1* ($p = 7.14 \times 10^{-04}$)

Figure S3: Top four significant genes showing differential circadian patterns from the human sleep restriction data. The two experimental conditions are the sufficient sleep condition and the insufficient sleep condition.
